# Neither the availability of D2 nor CP43 limits the biogenesis of PSII in tobacco

**DOI:** 10.1101/2020.08.31.272526

**Authors:** Han-Yi Fu, Rabea Ghandour, Stephanie Ruf, Reimo Zoschke, Ralph Bock, Mark Aurel Schöttler

## Abstract

The pathway of photosystem II assembly is well understood and multiple auxiliary proteins supporting it have been identified. By contrast, little is known about rate-limiting steps controlling PSII biogenesis. In the green alga *Chlamydomonas reinhardtii*, biosynthesis of the chloroplast-encoded D2 reaction center subunit (PsbD) limits PSII accumulation. To determine the importance of D2 synthesis for PSII accumulation in vascular plants and elucidate the contributions of transcriptional and translational regulation, the 5’-untranslated region of *psbD* was modified via chloroplast transformation in tobacco. A drastic reduction in *psbD* mRNA abundance resulted in a strong decrease of PSII content, impaired photosynthetic electron transport, and retarded growth under autotrophic conditions. Overexpression of the *psbD* mRNA also increased transcript abundance of *psbC* (the CP43 inner antenna protein), which is co-transcribed with *psbD*. Because translation efficiency remained unaltered, translation output of *pbsD* and *psbC* increased with mRNA abundance. However, this did not result in increased PSII accumulation. The introduction of point mutations into the Shine-Dalgarno-like sequence or start codon of *psbD* decreased translation efficiency without causing pronounced effects on PSII accumulation and function. These data show that neither transcription nor translation of *psbD* and *psbC* are rate-limiting for PSII biogenesis in vascular plants, and that PSII assembly and accumulation in tobacco are controlled by different mechanisms than in *Chlamydomonas*.

**One sentence summary:** PSII biogenesis in tobacco is neither limited by transcript accumulation nor translation of *psbD* and *psbC*.

## Introduction

Photosystem II (PSII), the water-plastoquinone oxidoreductase protein supercomplex of oxygenic photosynthesis, catalyzes the first step of linear electron flux in the thylakoid membranes of cyanobacteria and photosynthetic eukaryotes (Shen, 2015). Electron transfer within PSII is initiated by a light-induced charge separation at the reaction center (RC) chlorophyll-a dimer P_680_, which then transfers one electron to the first quinone acceptor Q_A_. From Q_A_, the electron is transferred to the plastoquinone-binding site (Q_B_-site) and reduces plastoquinone to plastosemiquinone. Following a second charge separation, the subsequent reduction of the plastosemiquinone to plastoquinol is coupled to the uptake of two protons from the stroma. Plastoquinol is released into the thylakoid membrane and re-oxidized at the cytochrome b_6_f complex (cyt b_6_f). After each charge separation, P_680_^+^ is reduced by the oxygen-evolving Mn_4_O_5_Ca complex (OEC) near the lumenal surface of PSII. After four oxidation steps, two water molecules are oxidized to molecular oxygen, and the Mn_4_O_5_Ca cluster is reduced again by the four electrons abstracted from the two water molecules (Dau et al., 2012; Vinyard and Brudvig, 2017).

PSII functions as a dimer, and each monomer is composed of more than 20 core subunits, and additional peripheral antenna proteins of the light-harvesting complex II (LHCII) type. With a molecular mass of up to 1300 kDa, the PSII-LHCII supercomplexes are the largest complexes of the photosynthetic apparatus (Dekker and Boekema, 2005; Kouril et al., 2012; Shen, 2015). The PSII RC core is formed by the D1 and D2 heterodimer that binds all redox-active cofactors necessary for rapid electron transfer from water to plastoquinone. D1 and D2 are encoded in the chloroplast genome (plastome) by the *psbA* and *psbD* genes, respectively. An additional redox-active cofactor, the heme of cytochrome b_559_ (cyt b_559_), is bound by PsbE and PsbF, which are also plastome-encoded. Cyt b_559_ is an essential structural component and required for PSII assembly, but its physiological function is enigmatic. It has been suggested to mediate a cyclic electron flux within PSII when the PSII donor side is inactive (Shinopoulos and Brudvig, 2012; Takagi et al., 2019), and may function as a plastoquinol oxidase (Bondarava et al., 2003; Bondarava et al., 2010).

The PSII RC is surrounded by the inner antenna proteins CP47 (PsbB) and CP43 (PsbC) also encoded in the plastome, and multiple membrane-intrinsic low molecular mass subunits encoded either in the plastome or the nucleus. Some of these subunits are essential for PSII accumulation or function, while others are not (reviewed by Shi et al., 2012; Plöchinger et al., 2016). Additionally, the three nuclear-encoded extrinsic subunits PSBO, PSBP, and PSBQ are associated with the lumenal side of PSII and stabilize the OEC (Bricker et al., 2012).

The process of *de novo* PSII assembly is largely conserved from cyanobacteria to vascular plants, except that the subcellular localization of some steps varies (Komenda et al., 2012; Nickelsen and Rengstl, 2013). Assembly proceeds in a modular fashion and starts with cyt b_559_, which stably accumulates even in the absence of the other PSII RC subunits (Müller and Eichacker, 1999; Kanervo et al., 2008; Plöscher et al., 2009; Schmitz et al., 2012). Subsequently, D2 is co-translationally inserted into the thylakoid membrane (Zoschke and Barkan, 2015) and binds to cyt b_559_, forming the D2-cyt b_559_ subcomplex (Komenda et al., 2004). The addition of a complex consisting of pre-D1, a D1 protein precursor with a short C-terminal extension, and PsbI leads to the formation of the “RC-like complex” (Dobakova et al., 2007), which is stabilized by multiple auxiliary proteins (Li et al., 2019). After maturation of the D1 protein by the lumenal C-terminal processing protease CTPA (Che et al., 2013), the “RC47 subcomplex” is formed by binding of CP47 (PsbB) and the rapid addition of PsbH, PsbR, and PsbTc (Rokka et al., 2005). Finally, CP43 (PsbC), PsbK, and PsbZ bind to the RC (Rokka et al., 2005; Boehm et al., 2011). This complex migrates into the grana stacks, where it is photoactivated by binding the extrinsic OEC subunits (Mamedov et al., 2000; Mamedov et al., 2008). Ultimately, two PSII monomers form a PSII dimer and bind additional LHCIIs (Shevela et al., 2016). During the assembly process, more than 20 auxiliary proteins located in the stroma, the thylakoids, and the lumen transiently bind to PSII. They protect and stabilize all assembly intermediates except for the early D2-cyt b_559_ subcomplex, which appears to accumulate without the support of auxiliary proteins (Shi et al., 2012; Nickelsen and Rengstl, 2013; Plöchinger et al., 2016). Some auxiliary proteins mediate chlorophyll and cofactor insertion into the nascent complex (Hey and Grimm, 2020).

In vascular plants, PSII contents are highly variable (reviewed by Schöttler and Toth, 2014). In *Arabidopsis thaliana*, PSII contents increase almost fourfold from 2.5 to 9 mmol PSII per mol chlorophyll, at the expense of the peripheral LHCII, when the actinic light intensity is increased from 35 to 600 μE m^-2^ s^-1^ (Bailey et al., 2001). Similar changes have been observed in tobacco (*Nicotiana tabacum*; Petersen et al., 2011; Schöttler et al., 2017) and in *Chamerion angustifolium* (Murchie and Horton, 1998). The mechanisms, by which these adjustments are achieved, are largely unknown.

In principle, PSII accumulation could be limited by the biosynthesis of one of its subunits, by cofactor synthesis and insertion into the nascent complex, or via regulation of the assembly process itself, for example, by changes in the abundance of auxiliary proteins. While PSII assembly starts with cyt b_559_, a limitation by cyt b_559_ seems unlikely. Free cyt b_559_ accumulates not only in etioplasts (Müller and Eichacker, 1998; Kanervo et al., 2008), but also in chloroplasts with severely disturbed sugar-phosphate metabolism and redox poise (Schmitz et al., 2012). Because D2 is next incorporated into the nascent PSII complex, its synthesis might control PSII accumulation. Indeed, in the green alga *Chlamydomonas reinhardtii*, D2 controls the accumulation of the other plastid-encoded subunits by a regulatory mechanism called “Control by Epistasy of Synthesis” (CES). The presence of D2 (or, more likely, of the D2-cyt b_559_ subcomplex) is required for efficient translation of D1, because unassembled D1 inhibits its own translation. Likewise, the presence of the “RC-like complex” as acceptor for newly translated CP47 is a prerequisite for the translation of *psbB*, as again, unassembled CP47 blocks its own synthesis (Minai et al., 2006). While cyt b_559_ is also essential for PSII accumulation in *C. reinhardtii* (Morais et al., 1998), it does not control synthesis of D2.

In *C. reinhardtii, psbD* mRNA abundance directly restricts PSII accumulation. A point mutation in the *psbD* promoter reduces both *psbD* mRNA accumulation and D2 protein content to 35% of the wild-type level (Klinkert et al., 2005). Furthermore, because the AUG start codon is part of an mRNA secondary structure, *psbD* cannot be spontaneously translated (Klinkert et al., 2006). Mutations of this repressor element not only resulted in increased translation of the *psbD* mRNA, but also led to a 20% increase in PSII accumulation, again supporting a limiting role of PsbD in PSII biogenesis (Klinkert et al., 2006). In the wild type, translation initiation of *psbD* is controlled by the translational activator RBP40 and the RNA stabilization factor Nac2, which open up the mRNA secondary structure and thereby facilitate translation initiation (Schwarz et al., 2007). However, neither RBP40 nor Nac2 have orthologs in vascular plants, and while *psbD* is encoded by a monocistronic mRNA in *C. reinhardtii*, in vascular plant plastomes, it forms an operon with*psbC* (Yao et al., 1989; Christopher et al., 1992; Adachi et al., 2012). In tobacco, the main promoter of this operon is located 905 bp upstream from the ATG codon of *psbD*. Additionally, a promoter located 194 bp upstream from the start codon of *psbC* is located within the protein-coding region of *psbD* (Yao et al., 1989). Because the reading frames of *psbD* and *psbC* overlap by 17 nucleotides in tobacco, it has been suggested that their translation is partly coupled. At least some of the ribosomes released after termination of *psbD* translation may immediately rebind to the *psbC* 5’-untranslated region (UTR) and initiate CP43 synthesis (Adachi et al., 2012). However, because *psbC* can be also expressed as a monocistronic transcript (Yao et al., 1989), translational coupling is no prerequisite for PsbC synthesis.

While in higher plant chloroplasts, translation regulation is usually considered to be more important than transcript abundance (reviewed by Zoschke and Bock, 2018), a limitation of photosynthetic complex biogenesis by transcript abundance was recently reported for the PetA subunit of the cyt b_6_f in tobacco (Schöttler et al., 2017). To determine if D2 or CP43 plays a limiting role for PSII accumulation in vascular plants, here we have generated transplastomic tobacco mutants with altered *psbD-psbC* transcript abundances. To assess a potential role of *psbD* translation regulation, we altered *psbD* translation initiation by the introduction of point mutations into the ATG start codon. Similar approaches were used by Rott et al. (2011) to reduce *atpB* translation, and by Moreno et al. (2017) to repress translation of ClpP, the chloroplast-encoded catalytic subunit of the stromal Clp protease (Nishimura and van Wijk, 2015). Also, the Shine-Dalgarno-like sequence (SD) in the 5’-UTR of *psbD* was mutated. The SD interacts by complementary base pairing with the anti-SD sequence in the 16S ribosomal RNA to ensure proper positioning of the translation initiation complex close to the start codon. In a transplastomic tobacco line carrying a mutation in the anti-SD sequence, *psbD* ranked among the genes, whose translation was most severely affected (Scharff et al., 2017). Our data suggest that *psbD* transcript abundance is critical in controlling D2 synthesis, but D2 is not the rate-limiting subunit for the accumulation of PSII in tobacco. We also show that the regulation of D2 synthesis is markedly different between *C. reinhardtii* and vascular plants.

## Results

### Generation of transplastomic tobacco lines with altered *psbD* expression

To dissect the contributions of transcriptional and translational regulation to the synthesis of the plastid-encoded D2 subunit, we generated chloroplast mutants with altered transcription of the *psbD-psbC* operon by disrupting the *psbD* 5’-UTR via insertion of the selectable marker gene *aadA* (for details, see Methods). In one set of transplastomic mutants, the selectable marker, whose expression is driven by the strong *Prrn* promoter, was inserted in the plastome in sense orientation relative to *psbD*. These mutants will subsequently be referred to as “sense” mutants (s-mutants). Read-through transcription from the *aadA* gene is expected to result in strong over-expression of the downstream *psbD-psbC* transcription unit (Zhou et al., 2007; Lu et al., 2013; Zhang et al., 2015). In another set of mutants, the selectable marker was inserted in antisense orientation relative to *psbD*, so that no read-through transcripts can occur. These mutants are referred to as “antisense” (as)-mutants. Because insertion of the selectable marker in antisense orientation may negatively affect the expression of neighbouring genes transcribed in the opposite direction (Loiacono et al., 2019), transcript accumulation of *psbD* could be strongly reduced in all as-mutants. Additionally, to alter translation initiation efficiency, we introduced point mutations into the translation initiation codon. The standard ATG translation initiation codon was replaced either by a GTG or TTG codon (**Figure 1A**), which are both recognized by the chloroplast ribosome as translation initiation sites, but with lower efficiency than the ATG codon (Rott et al., 2011; Moreno et al., 2017). Furthermore, as Scharff et al. (2017) had shown the SD dependency of *psbD* translation, we mutated the SD of *psbD* from GGAGGA to GGACGA, GGAGCA, or GGACCA.

**Figure 1.**
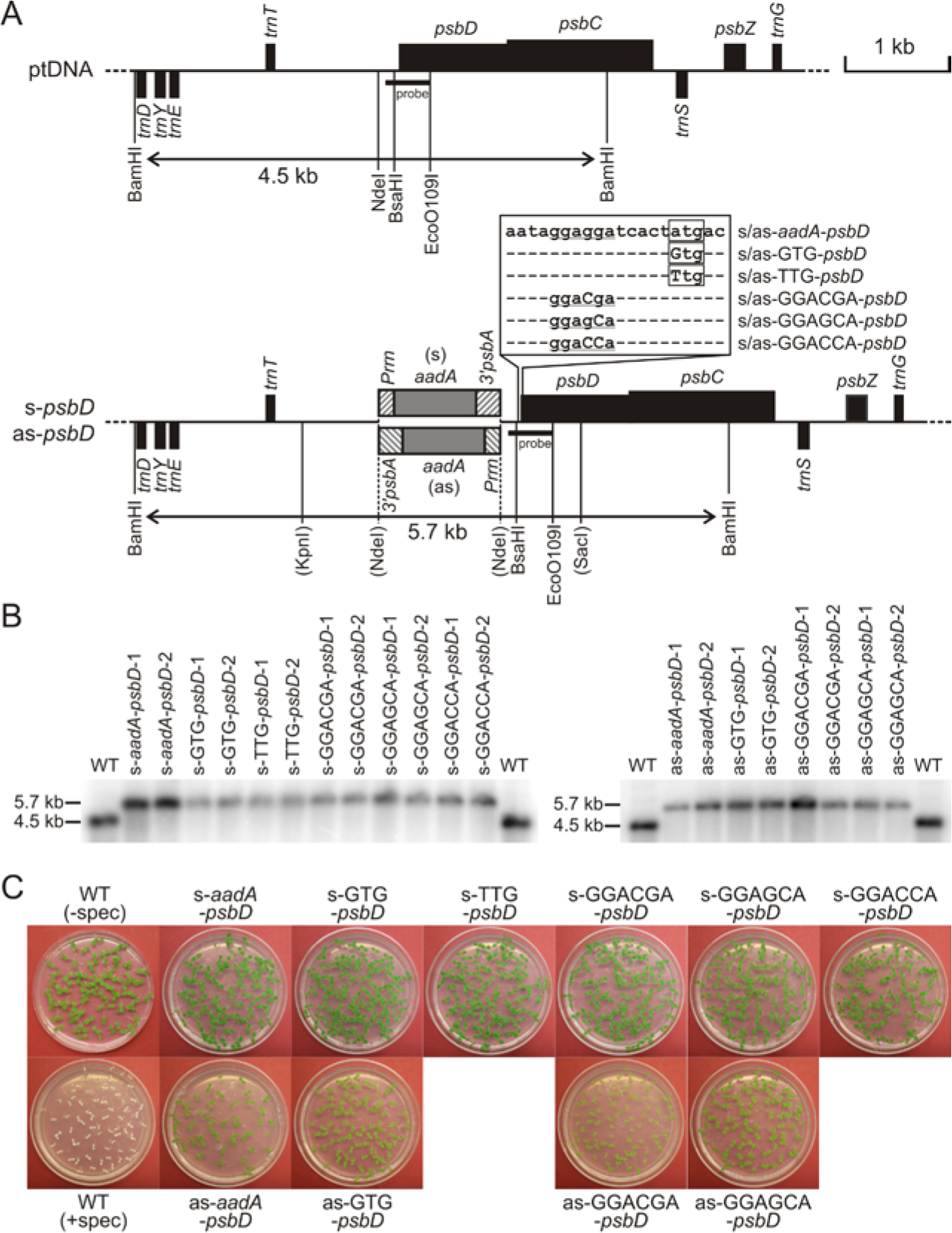
Generation and molecular characterization of *psbD* mutants. (A) Physical maps of the wild-type and mutant plastomes including all restriction sites used for cloning and RFLP analyses. Genes above the line are transcribed from left to right, genes below the line are transcribed in the opposite direction. The position of the probe used for RFLP analyses and the sizes of the detected BamHI restriction fragments are indicated. (B) RFLP analyses of transplastomic tobacco lines. DNA samples of transplastomic and wild-type (WT) plants were digested with the restriction enzyme BamHI, generating a 4.5 kb fragment in the wild type. Exclusive presence of the transplastomic fragment of 5.7 kb in all mutants indicates their homoplasmic state (left panel: sense-mutants, right panel: as-mutants). (C) Inheritance tests to confirm homoplasmy. Seeds from the wild type and the different sense and antisense mutants were germinated in the presence of spectinomycin.

Transplastomic lines were obtained by particle-gun mediated chloroplast transformation and selection for spectinomycin resistance (conferred by the *aadA* gene). All mutants could be generated and propagated in tissue culture under mixotrophic conditions. However, when plants were transferred to autotrophic growth conditions, neither the as-TTG-*psbD* mutant plants nor the as-GGACCA-*psbD* mutant plants survived. Both mutants could only be maintained in tissue culture, and consequently, did not produce seeds. Therefore, these mutants were excluded from further analysis. The homoplasmic state of all other mutants was tested by restriction fragment length polymorphism (RFLP) analysis using a *psbD*-specific probe and BamHI as restriction enzyme (as indicated in the map in **Figure 1A)**. Two representative independent lines for each construct were analyzed (**Figure 1B**). While the wild type showed the expected restriction fragment of 4.5 kb size, all transplastomic mutants showed exclusively a larger fragment of 5.7 kb, which arises from the insertion of the *aadA* marker gene with its promoter and terminator. The presence of the point mutations was confirmed by DNA sequencing.

Absence of the smaller wild-type-like fragment in the mutants strongly suggested the homoplasmic state of all transplastomic lines. To ultimately confirm homoplasmy, we also performed seed germination tests on spectinomycin-containing medium. This is the most sensitive assay to distinguish between homoplasmic and heteroplasmic transformants (Svab and Maliga, 1993; Bock, 2001; Bock 2015). As expected, cotyledons of wild-type seedlings germinated in the presence of spectinomycin were completely white, because inhibition of plastid translation prevents assembly of the photosynthetic apparatus (**Figure 1C**). By contrast, all transplastomic lines produced a homogeneous population of green seedlings, thus unequivocally confirming their homoplasmic state. Interestingly, seedlings from the as-mutants looked more light green compared to those from s-mutants, which is in line with the phenotypes of as-mutants grown in soil (see below).

### Transcript accumulation from the *psbD* operon

To determine general effects of the insertion of the selectable marker gene and the introduction of the point mutations on mRNA accumulation and processing, northern blot analysis was performed for all s-mutants, the as-*aadA-psbD* control line, and the as-GTG-*psbD* and as-GGAGCA-*psbD* mutants, which were viable under autotrophic conditions (**Figure 2**). Even though the as-GGACGA-*psbD* plants had produced some seeds (and, therefore, could be included in the germination assays; **Figure 1C**), they only rarely survived autotrophically. Therefore, this mutant was excluded from further analyses. In the wild type, using a *psbD*-specific probe, a complex pattern of transcripts between 4.4 and 2.6 kb size was observed. It arises from mRNA cleavage at a processing site located at position −132 relative to the translation initiation codon (Adachi et al., 2012), and from transcriptional read-through into *psbZ*. The *psbZ* gene is encoded downstream of *psbC* on the same DNA strand (**Figures 1A and 2B**).

The mutants with *aadA* inserted in s-orientation revealed the presence of two novel bands, which represent read-through transcripts from the strong *Prrn* promoter in front of the *aadA* selectable marker gene through the *psbD-psbC* oper

**Figure 2.**
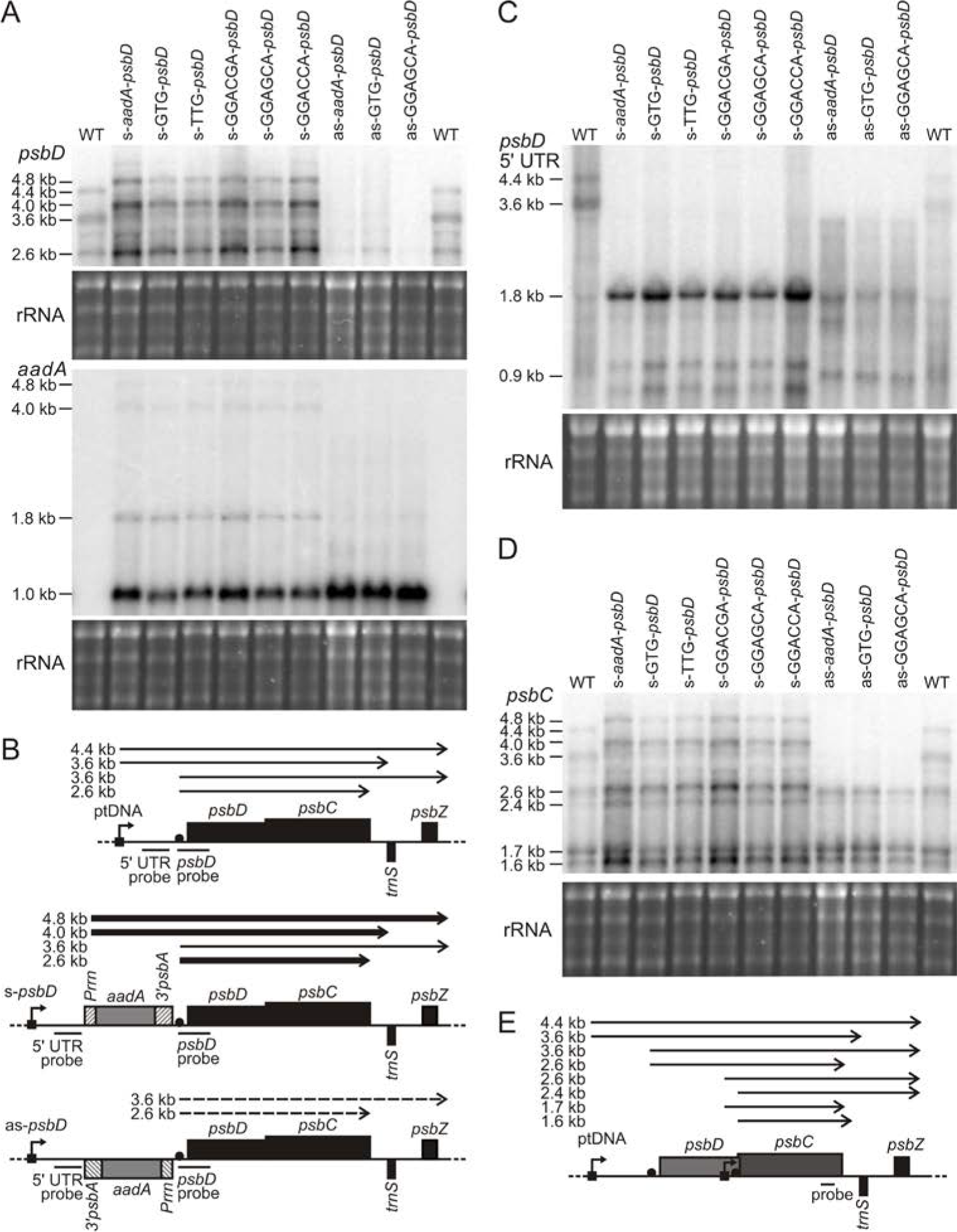
Changes in *psbD* and *psbC* transcript patterns and transcript abundance in transplastomic sense and antisense mutants. (A) Northern blots probed with *psbD*-(upper panel) and *aadA*-specific (lower panel) probes. As a loading control, the ethidium bromide-stained agarose gel prior to blotting is shown below each blot. WT: wild type. (B) Schematic representation of the *psbD* operon and the transcript species giving rise to the bands observed in northern blots. (C) Northern blot hybridized to a probe against the 5’-UTR of *psbD*. (D) Probe against *psbC*. (E) Schematic representation of the *psbC*-containing transcripts detected by northern blot hybridization.

on (transcript of approximately 4.0 kb size), or even through *psbZ* (approximately 4.8 kb size). Furthermore, the abundance of the processed dicistronic mature *psbD-psbC* transcript of 2.6 kb size was strongly increased, suggesting that the presence of the *aadA* selectable marker does not interfere with the normal processing of *psbD-psbC* polycistronic transcripts. In the case of the as-lines, a drastic decrease in *psbD* transcript abundance was observed, indicating that the *aadA* insertion indeed massively reduced transcription from the *psbD* promoter. Only small amounts of the mature dicistronic transcript of 2.6 kb were detectable.

To confirm the identity of the novel large transcripts in the mutants, we used an *aadA*-specific probe (**Figure 2A**). As expected, it did not give any signal in the wild type, but in the s-mutants revealed bands at 4.8 and 4.0 kb size, which correspond to the mRNA species already detected with the *psbD* probe. A stronger band of 1.8 kb size likely represents the processing product cleaved in front of the *psbD* translation initiation site and still includes the major part of the 5’-UTR of the *psbD* transcript. The strong 1.0 kb signal only covers the *aadA* selectable marker gene terminated by the 3’-*psbA* element downstream of the *aadA* coding region. This band was also the only prominent signal in the as-mutants.

Next, we used a probe specific for the long 5’-UTR of the primary *psbD* transcript (**Figure 2C**). As expected, in the wild type, this probe allowed us to detect major bands of 4.4 and 3.6 kb size (as reported previously; Yao et al., 1989), likely corresponding to the full-length transcripts covering either only the *psbD-psbC* coding regions or additionally including *psbZ*. In the s-mutants, by far the most prominent band was a band of 1.8 kb size, which likely corresponds to the processed *aadA* mRNA transcribed from the *psbD* operon promoter. Finally, we also used a *psbC* probe (**Figure 2D**), which resulted in the most complex transcript pattern with more than six distinct bands (**Figure 2E**). In the s-mutants, increased accumulation especially of the full-length transcripts spanning from the selectable marker to *psbZ* and of the processed mRNAs was observed. In the as-mutants, the long transcripts starting from the promoter in front of *psbD* were undetectable. However, shorter transcripts of up to 2.6 kb length starting from the *psbC*-specific promoter within the *psbD* coding region (Yao et al., 1989) accumulated to almost normal levels. These either spanned both *psbC* and *psbZ*, due to incomplete transcription termination at the end of *psbC*, or covered only *psbC*.

### Chloroplast translation in the mutants

To obtain information on changes in *psbD* translation due to the changes in mRNA abundance and the point mutations introduced into the 5’-UTR or the start codon, we performed polysome loading analyses as a proxy for translation activity (Barkan, 1998). Material was harvested from young leaves with still active thylakoid biogenesis. mRNAs with different ribosome loading were separated by sucrose density gradient centrifugation into 12 fractions, with fraction 12 being the fraction of highest density (**Figure 3**). To determine the distribution of free, untranslated mRNAs, a puromycin-treated wild-type sample was also analyzed. Puromycin dissociates ribosomes from mRNAs. In the puromycin control, the *psbD-psbC* mRNA was mainly found in fractions three to five, while in the untreated wild type, mRNA distribution peaked in fractions seven to nine. Despite the massive increase in mRNA abundance in the s-*aadA-psbD* mutant, mRNA distribution and therefore translation efficiency per mRNA was unaltered, indicating increased total synthesis of PsbD. Both the s-GTG-*psbD* and s-TTG-*psbD* mutants showed a marked shift in mRNA distribution towards fractions of lower density. Point mutations in the SD also shifted mRNA distribution in the polysome profile upward by one to three fractions, and this effect was most pronounced in the s-GGACGA-*psbD* mutant. While the as-*aadA-psbD* mutant showed a polysome distribution very similar to the wild type, the as-GTG-*psbD* mutant and the as-GGAGCA-*psbD* mutant showed mild shifts in their polysome profiles by one to two fractions towards a lower density.

**Figure 3.**
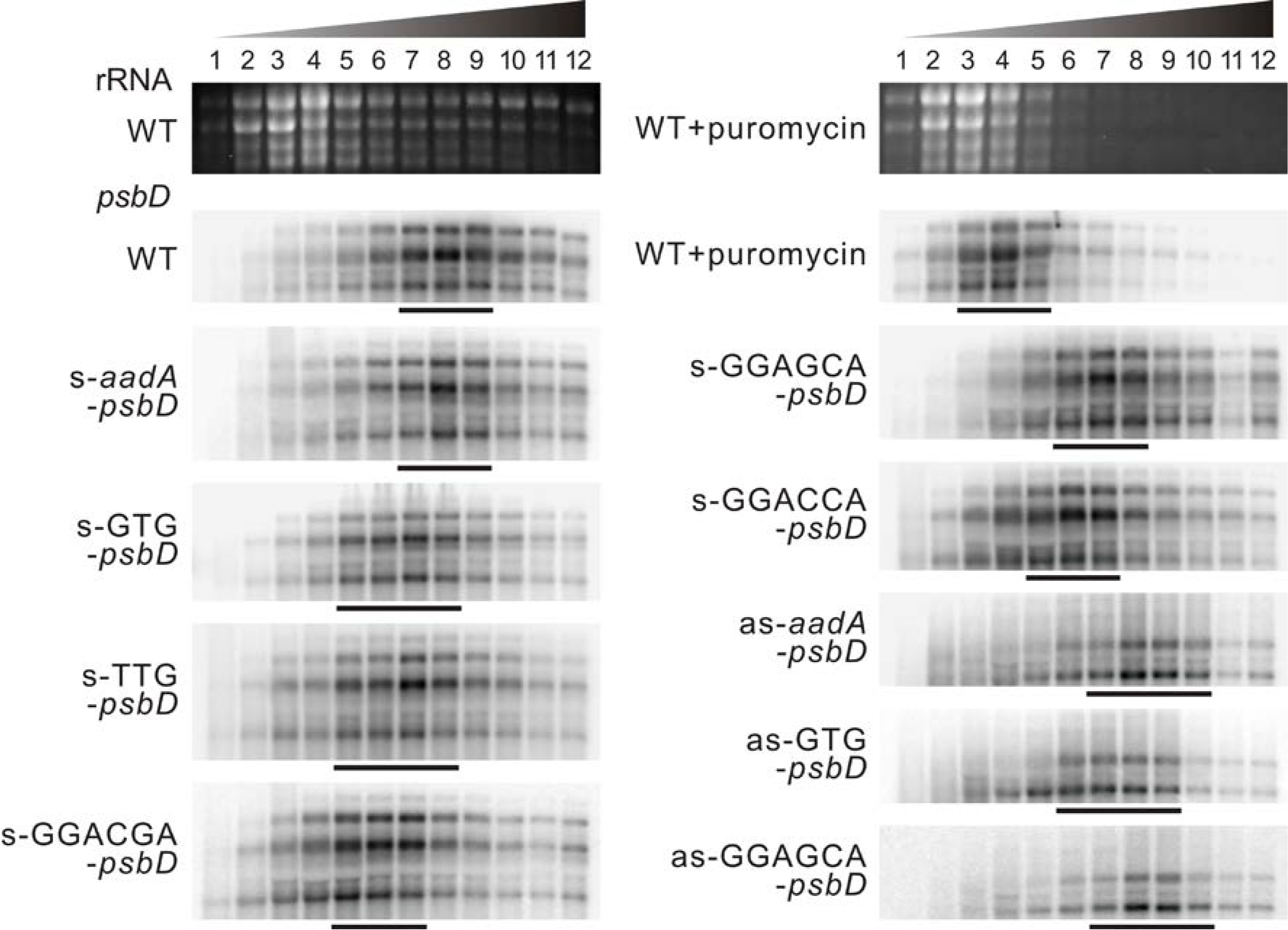
*psbD* polysome loading analyses of sense and antisense transformants grown on soil. Material was harvested from young leaves with still active thylakoid biogenesis. mRNAs with different ribosome coverage were separated by sucrose density gradient centrifugation into 12 fractions, with fraction 12 being the fraction of highest density. To determine the distribution of free, untranslated mRNAs, a puromycin-treated wild-type sample was also analyzed. Puromycin dissociates ribosomes from the mRNAs, as confirmed by comparison of ethidium bromide-stained agarose gels (upper panels). The major *psbD*-containing fractions are indicated by horizontal bars below each blot. WT: wild type.

Resolution and sensitivity of polysome analyses are rather limited, and in the case of polycistronic transcripts, due to the physical linkage of the reading frames, polysome loading only provides an average value of the total translation of the entire transcript (reviewed by Zoschke and Bock, 2018). Therefore, translation of *psbD* and *psbC* cannot be distinguished, and the altered polysome profiles could be due to altered translation of either *psbD* or*psbC*, or both. For example, *psbD* translation might alter translation of *psbC* either due to translational coupling (Adachi et al., 2012), or some other kind of feedback mechanism (such as CES in *C. reinhardtii*; Minai et al., 2006).

To characterize transcription and translation in more detail, we selected the s-*aadA-psbD* control mutant, the two s-translation initiation codon mutants, and the wild type for a comprehensive analysis of changes in transcript accumulation and translation of all plastome-encoded genes by transcript and ribosome profiling. To this end, transcript abundances and ribosome footprints were analyzed using high-resolution tiling arrays and then averaged for each chloroplast open reading frame as previously described (Trösch et al. 2018). In each case, ribosomal footprints (“translation output”) and transcript abundances (“RNA”) of the three different mutants were plotted against the signals of wild-type tobacco analyzed in the same experiment (**Figure 4,** x-axis). To determine the relative translation efficiency of each gene, data were log_2_-transformed and transcript abundance was subtracted from ribosomal footprints (**Supplementary Figure 1;** Zoschke et al., 2013). The data shown represent averages from 3 biological replicates (see **Supplementary Figure 2**). Pronounced changes (≥ 2-fold) are highlighted in red color in **Figure 4**.

**Figure 4.**
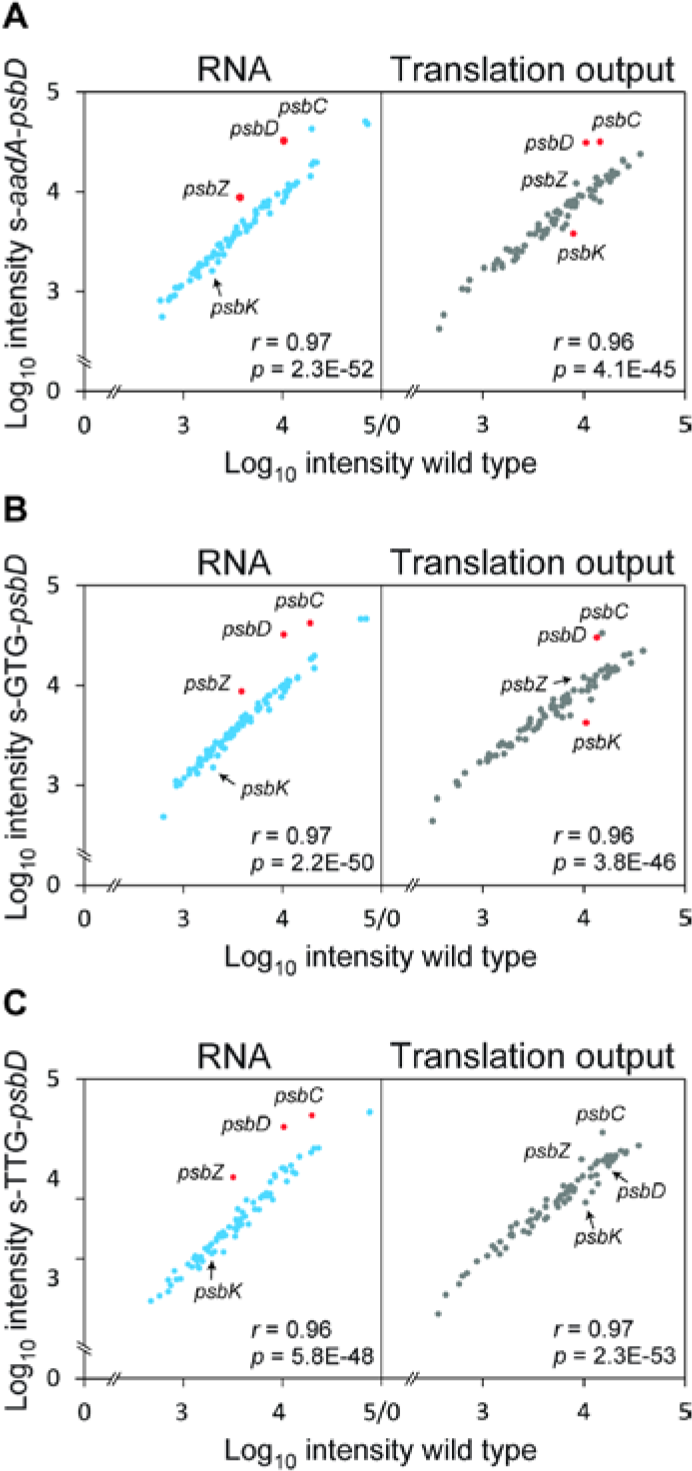
Relative translation output and RNA accumulation levels of transplastomic *psbD* mutant lines. Relative transcript accumulation (RNA) levels and chloroplast ribosome footprint (translation output) were examined by microarray-based ribosome profiling and compared between mutant and wild type. Average ribosome footprint and transcript abundances were calculated for each reading frame as described in Methods, and the mean of three biological replicates was log_10_-transformed and plotted for the transplastomic lines s-*aadA-psbD* (**A**), s-*psbD*-GTG (**B**) and s-*psbD*-TTG (**C)** against the corresponding wild type (left panels: transcript levels, right panels: ribosome footprints; for better visualization x- and y-axes are broken). Genes showing more than 2-fold changes are highlighted with red data points and named beside the data points in all analyses. Pearson’s r and Anova’s p-value (in nEm non-superscript format for n•10m) are given within each plot.

When the wild type was compared to the s-*aadA-psbD* control mutant (**Figure 4A**), we observed strong increases in transcript abundance of *psbD* and *psbZ*, in line with the changes observed by northern blot analysis (**Figure 2**). *psbC* transcript abundance was also increased, but remained slightly below twofold. Abundance of all other chloroplast transcripts did not change substantially. The translation output of *psbD* and *psbC*, but not of *psbZ*, was significantly increased. Unexpectedly, translation of *psbK* was clearly decreased. However, at the level of the relative translation efficiency, not a single significant difference between wild type and the s-*aadA-psbD* control mutant was observed (**Supplemental Figure 1**). The increased transcript abundance of *psbD* and *psbC* directly resulted in increased translation output, in agreement with their wild-type like polysome distribution profiles (**Figure 3**).

In the s-GTG-*psbD* mutant (**Figure 4B**), *psbD, psbC* and *psbZ* mRNA accumulation levels were more than twofold increased. This resulted in a pronounced increase in the translation output of both *psbD* and *psbC*, suggesting that the GUG initiation codon is almost as efficient as the standard AUG translation initiation codon for *psbD*. Similar to the other two s-mutants, the s-TTG-*psbD* mutant displayed significant increases in *psbD, psbC*, and *psbZ* transcript abundances (**Figure 4C**). However, different from the s-GTG-*psbD* mutant and despite its increased mRNA abundance, the s-TTG-*psbD* mutant did not show a significantly increased translation output for any of these reading frames. This is due to a strong reduction in the translation efficiency of especially *psbD*, indicating that the UUG initiation codon is much less efficient than AUG and GUG (**Supplemental Figure 1**). The strongly diminished translation efficiency of *psbD* did not virtually hamper translation of *psbC*. Consequently, *in vivo, psbC* translation can be considered as uncoupled (or only marginally coupled) to *psbD* translation initiation, contrasting a conclusion that was previously drawn based on *in vitro* translation data (Adachi et al., 2012).

### Only minor differences in photosynthetic parameters of s-mutants

To determine effects of the altered *psbD* transcript accumulation and translation initiation on plant growth and photosynthetic performance, the plants expressing the selectable marker gene in s-orientation were grown in soil under long-day conditions and with an actinic light intensity of 350 μE m^-2^ s^-1^. Plant growth and all physiological parameters analyzed (see below) were not significantly different between independent mutant lines harboring the same construct. Therefore, the growth phenotype of one representative plant per construct (**Figure 5A**), and average values of all mutant lines per construct (**Table 1, Figure 7**) are shown. Growth of all mutants was only slightly retarded relative to that of the wild type (**Figure 5A**).

**Table 1:** Chlorophyll content, chlorophyll a/b ratio, leaf absorptance, maximum quantum efficiency of PSII in the dark-adapted state (F_V_/F_M_), and photosynthetic complex contents per leaf area of sense-*psbD* mutants grown at an actinic light intensity of 350 μE m^-2^ s^-1^. Red color highlights values, which are significantly different from wild type.

| Parameter | WT | s-<br>*aadA-*<br>*psbD* | s-<br>*GTG-*<br>*psbD* | s-<br>*TTG-*<br>*psbD* | s-<br>*GGACGA-*<br>*psbD* | s-<br>*GGAGCA-*<br>*psbD* | s-<br>*GGACCA-*<br>*psbD* |
| --- | --- | --- | --- | --- | --- | --- | --- |
| Chlorophyll<br>[mg / m<sup>2</sup>] | 442.3<br>± 98.4 | 454.7<br>± 102.5 | 436.1<br>± 74.6 | 364.7<br>± 66.1 | 520.9<br>± 91.1 | 519.3<br>± 72.1 | 492.1<br>± 42.1 |
| Chlorophyll<br>*a/b* | 4.04<br>± 0.16 | 3.84<br>± 0.07 | 3.80<br>± 0.11 | 3.72<br>± 0.13 | 3.85<br>± 0.11 | 3.85<br>± 0.19 | 3.82<br>± 0.10 |
| Leaf<br>absorptance<br>(%) | 88.5<br>± 1.8 | 88.2<br>± 2.2 | 88.4<br>± 2.1 | 86.9<br>± 1.9 | 89.8<br>± 1.2 | 89.6<br>± 1.2 | 89.3<br>± 0.6 |
| $F_v/F_m$ | 0.81<br>± 0.02 | 0.80<br>± 0.02 | 0.80<br>± 0.02 | 0.79<br>± 0.01 | 0.80<br>± 0.02 | 0.81<br>± 0.01 | 0.80<br>± 0.02 |
| PSII<br>[μmol / m<sup>2</sup>] | 1.11 ±<br>0.25 | 1.16 ±<br>0.21 | 1.06 ±<br>0.17 | 0.81 ±<br>0.20 | 1.14<br>± 0.17 | 1.20<br>± 0.19 | 1.05<br>± 0.12 |
| Cyt b<sub>6</sub>f<br>[μmol / m<sup>2</sup>] | 0.49<br>± 0.14 | 0.45<br>± 0.07 | 0.45<br>± 0.12 | 0.37<br>± 0.10 | 0.58<br>± 0.12 | 0.51<br>± 0.10 | 0.52<br>± 0.08 |
| Pc<br>[μmol / m<sup>2</sup>] | 2.21<br>± 0.65 | 2.55<br>± 0.66 | 2.43<br>± 0.59 | 1.90<br>± 0.73 | 2.93<br>± 0.46 | 2.78<br>± 0.55 | 2.83<br>± 0.41 |
| PSI<br>[μmol / m<sup>2</sup>] | 0.97<br>± 0.24 | 0.99<br>± 0.22 | 0.95<br>± 0.16 | 0.81<br>± 0.19 | 1.15<br>± 0.17 | 1.13<br>± 0.18 | 1.08<br>± 0.10 |

**Figure 5.**
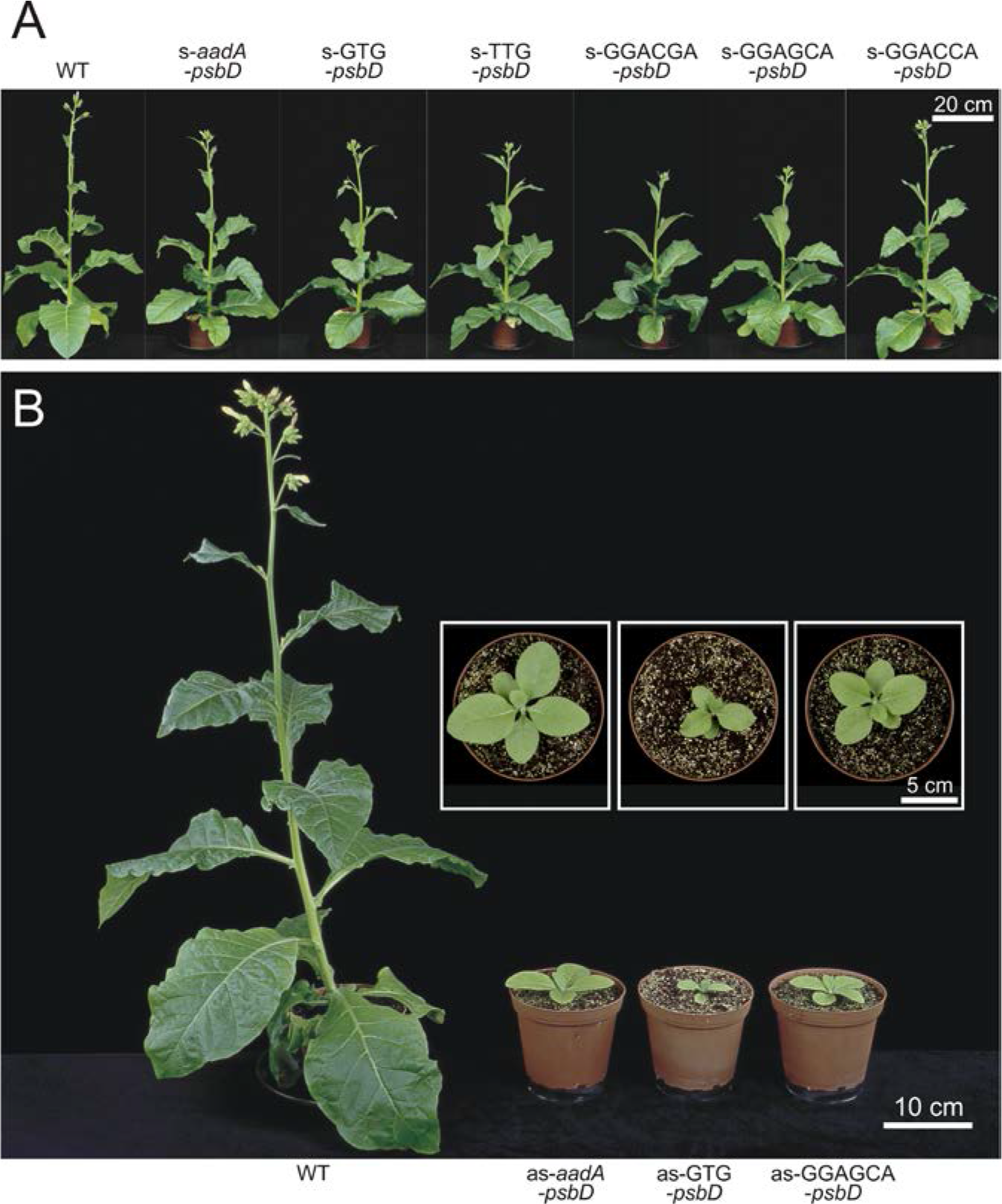
Growth phenotypes of transplastomic *psbD* mutants. (A) Growth phenotypes of the *s-psbD* transplastomic lines grown in soil at 350 μE m^-2^ s^-1^. The photographs were taken 10 weeks after seeds had been sown in soil, when the wild-type plants (WT) started to flower. (B) Growth phenotypes of the as-*psbD* transplastomic lines grown in soil at 100 μE m^-2^ s^-1^. The photographs were taken when the wild-type plants (WT) began to flower. The insets show close-up top views of the transplastomic plants.

To determine possible changes in the composition and function of the photosynthetic apparatus in detail, the chlorophyll a/b ratio, the chlorophyll content per leaf area, leaf absorptance, and the maximum quantum efficiency of PSII in the dark-adapted state (F_V_/F_M_) were determined for the youngest fully expanded leaves of plants at the onset of flowering (**Table 1**). Then, thylakoids were isolated from these leaves and the contents of both photosystems, the cyt b_6_f and plastocyanin (Pc) were determined spectroscopically from chemically or light-induced difference absorbance signals of cyt b_559_ (PSII), cytochromes b6 and f (cyt b_6_f) and P700 (PSI; see Methods). Finally, these data were re-normalized to a leaf area basis (**Table 1**). These quantifications were validated by immunoblots against essential subunits of the different photosynthetic complexes (**Figure 6**).

**Figure 6.**
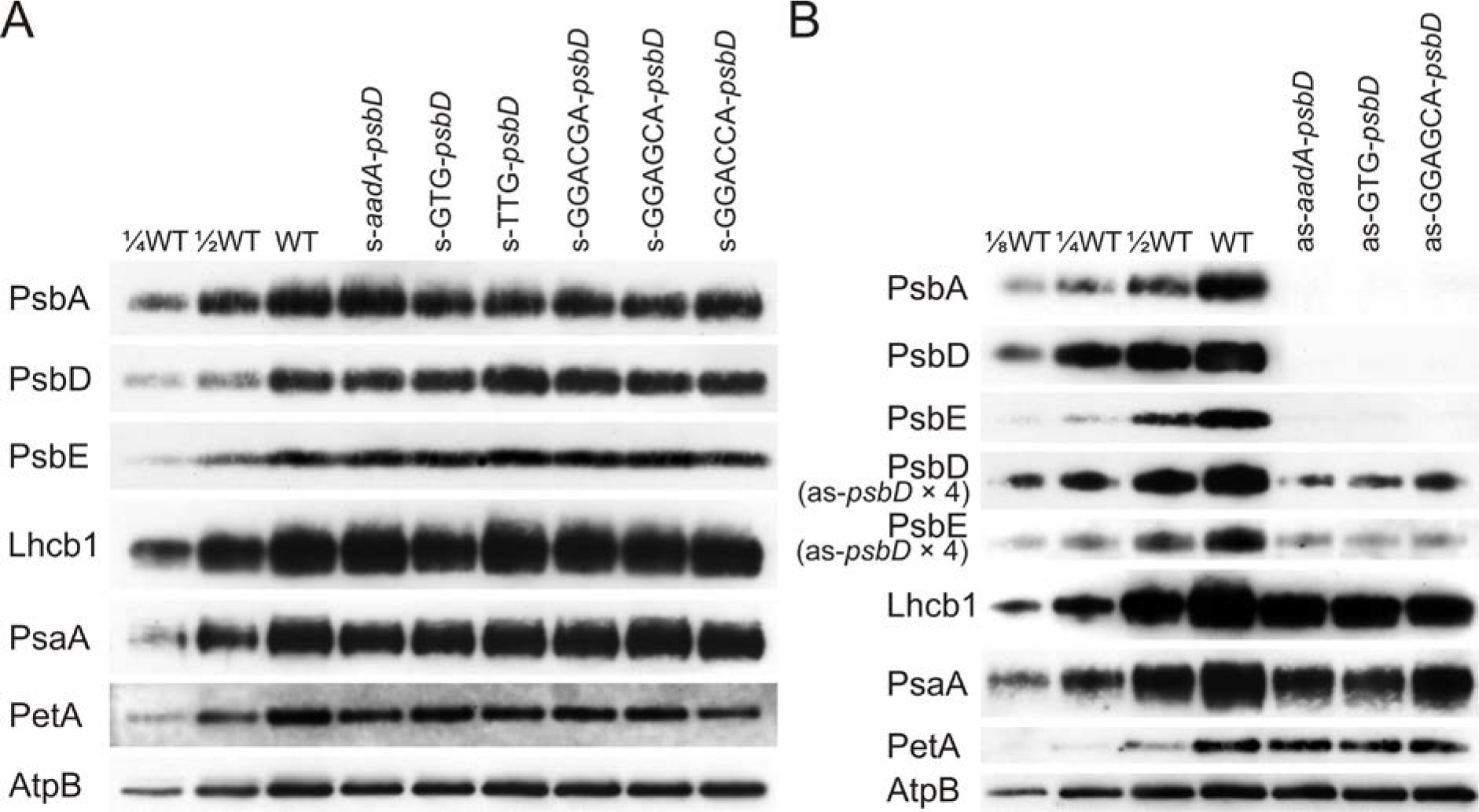
Protein accumulation in the *s-psbD* (A) and *as-psbD* (B) transplastomic lines. Immunoblot analyses were performed using antibodies against diagnostic subunits of photosystem II (PsbA, PsbD, and PsbE subunits), light-harvesting complex II (LHCB1), photosystem I (PsaA), the cytochrome b_6_f complex (PetA, i.e., cytochrome f), and the chloroplast ATP synthase (AtpB). Samples were loaded on an equal leaf area basis. For the wild type (WT), a dilution series is included to enable semiquantitative assessments. As accumulations of the PSII subunits were barely detectable in the as-*psbD* transplastomic lines (panel B), four times of the thylakoid amounts in these as-*psbD* lines (as-*psbD* × 4) were loaded along with unchanged amounts of WT samples.

Neither chlorophyll content nor leaf absorptance differed significantly between wild type and the s-mutants, while the chlorophyll a/b ratio of all mutants was slightly lower than in the wild type (**Table 1**). The maximum quantum efficiency of PSII in the dark-adapted state (F_V_/F_M_), as well as the PSII and cyt b_6_f content per leaf area, were only slightly lower in the s-TTG-*psbD* mutant than in the wild type. In these parameters, all other mutants did not differ from the wild type. For PSI accumulation, no significant differences could be observed for any of the s-mutants. These results were confirmed by immunoblots against the essential PSII RC core subunits PsbA (D1), PsbD (D2), and PsbE, a subunit of cyt b_559_. LHCB1 accumulation did not reveal any differences between the wild type and any of the s-mutants. When PSI, the cyt b_6_f, and chloroplast ATP synthase were probed with antibodies against their essential subunits PsaA, PetA (cytochrome f), and AtpB, no obvious differences in protein accumulation between mutants and wild type could be observed (**Figure 6**), in line with the spectroscopic results.

Accordingly, most mutants displayed only subtle changes in the light response curves of the chlorophyll-a fluorescence parameters qL (a measure for the redox state of the PSII acceptor side; Kramer et al., 2004) and qN (a measure for the thermal dissipation of excess excitation energy in the PSII antenna system, Krause and Weis, 1991). In the s-TTG-*psbD* mutant, the only s-mutant displaying a significant decrease in PSII accumulation, induction of qN was slightly shifted to lower light intensities, while for the reduction state of the PSII acceptor side, no significant differences were observed (**Figure 7A**). Finally, 77K chlorophyll-a fluorescence emission spectra were recorded and normalized to the PSI emission signal at 734 nm wavelength (**Figure 7C**). Again, no changes in the maximum emission signal of PSII at 686.5 nm wavelength were observable, strongly suggesting that the LHCII antenna proteins were well coupled to the RC, even in the s-TTG-*psbD* mutant.

**Figure 7.**
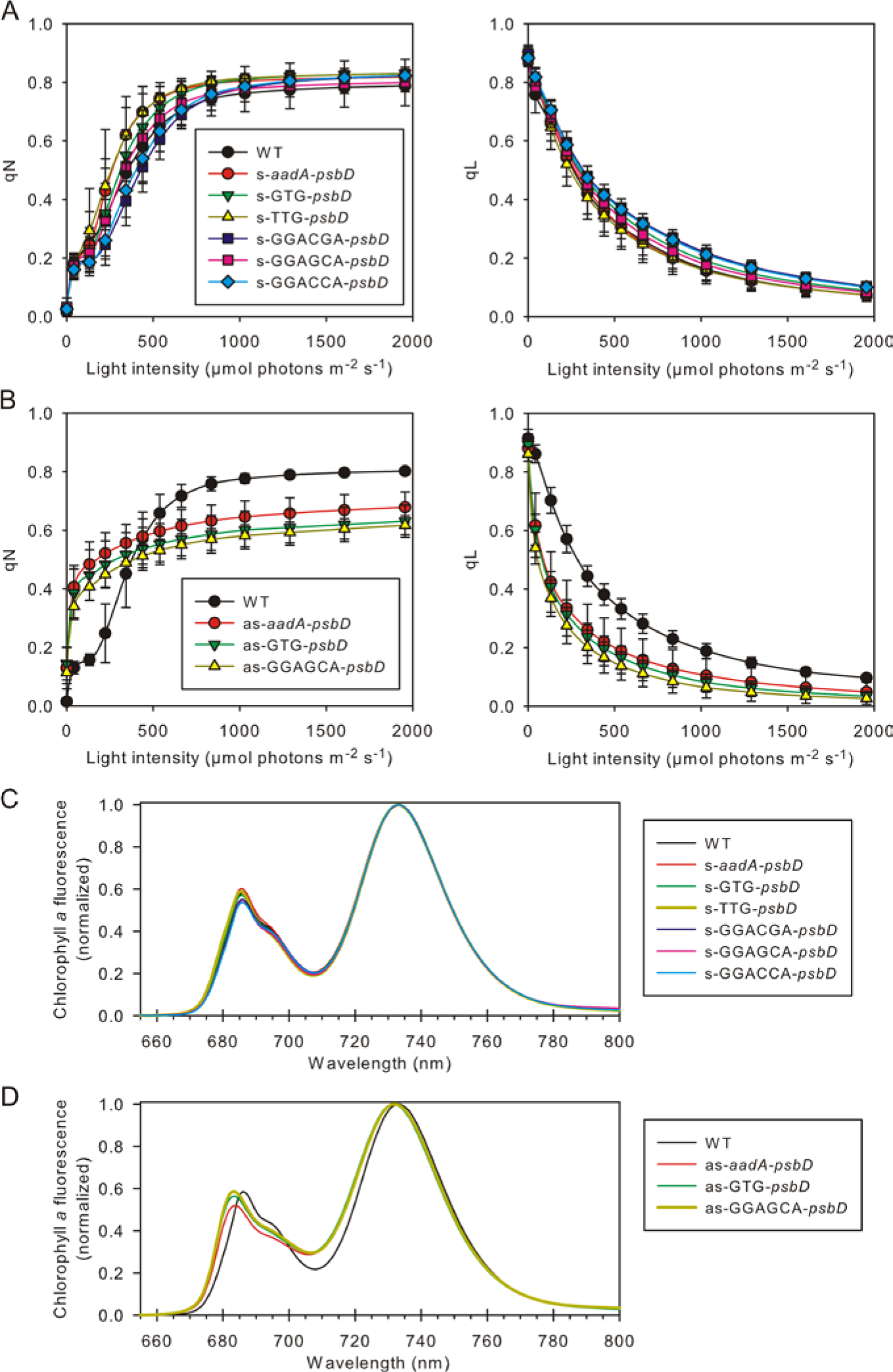
Chlorophyll-a fluorescence analysis of transplastomic *psbD* mutants. (A,B) Light response curves of qL (redox state of the PSII acceptor sides) and qN (non-photochemical quenching) in the s-*psbD* (A) and as-*psbD* (B) transplastomic lines. Error bars indicate the standard deviation of the mean, and the sample size is the same as indicated in Table 1. (C,D) 77K chlorophyll-a fluorescence emission spectra in the s-*psbD* (C) and as-*psbD* (D) transplastomic lines. The spectra are averaged per construct (n indicated in Table 1) and normalized to the PSI emission maximum at ~735 nm.

### as-mutants suffer from massive growth retardation and impaired photosynthesis

All three autotrophically viable as-mutants showed delayed growth and evidence of reduced photosynthetic performance. They were highly light-sensitive when grown at 350 μE m^-2^ s^-1^, the light intensity used for the characterization of the s-mutants. Therefore, the as-mutants were grown at a reduced light intensity of 100 μE m^-2^ s^-1^. Photographs in **Figure 5B** were taken when the wild type started to flower. At that time, all three mutants were strongly retarded in growth, most severely the as-GTG-*psbD* mutant. For a more detailed physiological analysis, to avoid artifacts due to differences in the developmental state of the plants, young fully expanded source leaves of non-flowering wild-type plants and mutants were used (**Table 2**). Despite the reduced growth light intensity, the data for wild-type tobacco are almost indistinguishable from those of the wild type grown at 350 μE m^-2^ s^-1^. This is unsurprising because, between 100 and 350 μE m^-2^ s^-1^, the photosynthetic apparatus of tobacco only shows minor responses to the growth light intensity (Schöttler et al., 2017).

**Table 2:** Chlorophyll content, chlorophyll a/b ratio, leaf absorptance, maximum quantum efficiency of PSII in the dark-adapted state (F_V_/F_M_), and photosynthetic complex contents per leaf area of *as-psbD* mutants grown at a reduced actinic light intensity of 100 μE m^-2^ s^-1^. Red color highlights values, which are significantly different from wild type.

| Parameter | WT | *as-aadA-psbD* | *as-GTG-psbD* | *as-GGAGCA-psbD* |
| --- | --- | --- | --- | --- |
| Chlorophyll [$\text{mg / m}^2$] | $413.9 \pm 45.6$ | $145.2 \pm 19.7$ | $128.5 \pm 11.7$ | $124.7 \pm 26.6$ |
| Chlorophyll *a/b* | $4.18 \pm 0.11$ | $3.79 \pm 0.15$ | $3.94 \pm 0.09$ | $3.83 \pm 0.09$ |
| Leaf absorptance (%) | $87.7 \pm 1.1$ | $73.0 \pm 2.8$ | $71.3 \pm 2.5$ | $70.4 \pm 3.3$ |
| $F_v/F_m$ | $0.80 \pm 0.01$ | $0.28 \pm 0.07$ | $0.25 \pm 0.02$ | $0.24 \pm 0.05$ |
| PSII [$\mu\text{mol / m}^2$] | $1.19 \pm 0.14$ | $0.22 \pm 0.05$ | $0.23 \pm 0.05$ | $0.19 \pm 0.02$ |
| Cyt $b_6f$ [$\mu\text{mol / m}^2$] | $0.51 \pm 0.11$ | $0.29 \pm 0.05$ | $0.24 \pm 0.10$ | $0.31 \pm 0.08$ |
| Pc [$\mu\text{mol / m}^2$] | $2.27 \pm 0.41$ | $1.24 \pm 0.32$ | $1.35 \pm 0.25$ | $1.19 \pm 0.19$ |
| PSI [$\mu\text{mol / m}^2$] | $1.01 \pm 0.11$ | $0.34 \pm 0.05$ | $0.32 \pm 0.03$ | $0.31 \pm 0.07$ |

All parameters tested were significantly different between the wild type and the three as-mutants. The chlorophyll content per leaf area in the as-mutants was reduced to less than 35% of the wild-type level. In line with the lower chlorophyll content, leaf absorptance decreased as well. The reduction in the chlorophyll a/b ratio was less drastic, indicating a global reduction of all components of the photosynthetic apparatus, instead of a selective loss of PSII. Specific impairments in PSII RC accumulation should result in a strong decrease of the chlorophyll a/b ratio, because RCs only bind chlorophyll a, while the antenna proteins of the photosystems bind both chlorophyll a and b. The maximum quantum efficiency of chlorophyll-a fluorescence was drastically reduced in all three as-mutants, suggesting PSII photoinhibition or the presence of uncoupled LHCII in the thylakoid membrane. In line with the strongly reduced chlorophyll content and the only moderately affected chlorophyll a/b ratio, the accumulation of all photosynthetic complexes per leaf area was strongly decreased as well: PSII contents decreased to less than 20% of wild-type levels in all three mutants, and also PSI accumulation was severely reduced to approximately 30% of wild-type levels. The pronounced effect of reduced PSII contents on PSI accumulation is unsurprising, in that similar findings have been reported for *A. thaliana* antisense mutants against the PSBO subunit of the OEC (Dwyer et al., 2012). The reduction in both photosystems was more pronounced than that in the total chlorophyll content, indicating that LHC accumulation was affected to a lesser extent. Also, the cyt b_6_f and Pc were less strongly affected than the photosystems, decreasing only by 40 to 50% in content, relative to the wild type. Changes in photosynthetic complex accumulation were again confirmed immunologically (**Figure 6B**). Again, as proxies for PSII accumulation, PsbA (D1), PsbD (D2), and PsbE were probed. Accumulation of all PSII subunits was found to be strongly decreased. In line with the other redox-active complexes being less affected than PSII, accumulation of PsaA (PSI) and PetA (cyt b_6_f) were less severely reduced than that of the PSII subunits.

On a functional level, the light response curves of qL and qN revealed pronounced differences between the wild type and the as-mutants. Both the induction of qN and the reduction of the PSII acceptor side were strongly shifted to lower light intensities, while the full induction of qN in saturating light was impaired in the as-mutants (**Figure 7B**). Finally, in the 77K chlorophyll-a fluorescence emission spectra, the maximum emission wavelengths of PSI-LHCI and especially of PSII-LHCII were clearly shifted to lower wavelengths, indicating the presence of free, uncoupled LHCII and LHCI antenna proteins (**Figure 7D**). Uncoupled LHCI emit fluorescence at 77K with maxima between 701 nm and 730 nm wavelength (Castelletti et al. 2003; Croce et al., 2004), while the coupled LHCI-PSI in the wild type show their typical emission maximum at 734 nm wavelength. Likewise, the shift of the PSII-LHCII emission maximum from 686.5 nm to 682 nm indicates the presence of free, uncoupled LHCII, whose emission maximum is at 680 nm wavelength (Krause and Weis, 1991).

## Discussion

The biogenesis of PSII has been studied for decades, and its assembly sequence is well established. Also, more than 20 auxiliary proteins stabilizing different assembly intermediates and supporting the insertion of pigments and co-factors have been identified (Nickelsen and Rengstl, 2013; Lu, 2016). By contrast, much less is known about the limiting steps of PSII biogenesis, and how PSII accumulation is adjusted to different growth light intensities, which can result in more than fourfold changes in PSII content (reviewed by Schöttler and Toth, 2014). Therefore, identifying the rate-limiting step(s) of PSII biogenesis is of major importance to better understand the acclimation of the photosynthetic apparatus to different environmental conditions, and might pave the way for targeted manipulations of PSII content, without altering its subunit composition and function.

Here, we have addressed a potential limiting role of the two plastome-encoded subunits D2 and CP43, which are expressed from a single operon-type transcription unit in tobacco. Our main focus was on D2, which together with cyt b_559_ initiates the PSII assembly process. We did not consider cyt b_559_ as a candidate for controlling PSII accumulation, because much higher amounts of cyt b_559_ than of functional PSII were shown to accumulate in *A. thaliana* mutants with massively disturbed primary metabolism (Schmitz et al., 2012). In *C. reinhardtii*, a reduction in *psbD* mRNA directly resulted in a proportional reduction in D2 protein and PSII abundance, pointing to an important regulatory role of *psbD* transcript accumulation (Klinkert et al., 2005). Mutations in a repressor element of translation initiation in the 5’-UTR of *psbD* increased not only *psbD* translation initiation efficiency, but also PSII accumulation by up to 20% (Klinkert et al., 2006). Likely, increased D2 availability activates the synthesis of D1 by rapidly binding newly translated D1, which then cannot repress its own translation (Minai, 2006). A similar overaccumulation of functional PSII by increased synthesis of D2 (either due to increased *psbD* mRNA abundance or increased rates of translation) would be the ultimate proof that, similar to the situation in *C. reinhardtii*, D2 is also limiting PSII biogenesis in vascular plants.

### Neither D2 nor CP43 limit PSII accumulation in tobacco

To generate transplastomic tobacco with altered *psbD* transcript abundance, we disrupted the *psbD* 5’-UTR via insertion of the selectable marker gene *aadA*. In the s-mutants, read-through transcription from the strong *Prrn* promoter of the *aadA* massively increased *psbD* (and *psbC*) transcript abundance (**Figure 2**). This increase in *psbD* mRNA levels was further confirmed by the chloroplast transcriptome profiling data (**Figure 4**), which additionally showed a strong increase in *psbZ* mRNA abundance. This again is a consequence of transcriptional read-throughs through the *psbD-psbC* dicistron into *psbZ* located further downstream. In the as-mutants, the disruption of the native promoter strongly decreased *psbD* transcript abundance. Therefore, the situation in the as-mutants is similar to the reduced *psbD* transcript accumulation in the *C. reinhardtii* promoter mutant (Klinkert et al., 2005). Also in tobacco, the massive decrease in transcript abundance resulted in a more than 80% reduction in PSII accumulation (**Table 2**), which could not be compensated by increased translation of the residual mRNAs (**Figure 3**). This observation is in line with a limiting role of D2 synthesis for PSII assembly.

The strong increase in *psbD* and *psbC* transcript in the s-*aadA-psbD* mutant did not result in increased PSII accumulation, even though mRNA distribution in the sucrose density gradients used for polysome analysis remained unchanged, indicating that the supernumerary mRNA molecules undergo translation (**Figure 3**). This conclusion is also supported by ribosome footprint analyses, which show proportional increases of both transcript abundance and translation output for *psbD* and *psbC* (**Figure 4A**). Consequently, relative translation efficiency remained unaltered (**Supplemental Figure 1**). However, neither D2 protein level (**Figure 6A**) nor PSII content (**Table 1**) increased. Also, all other photosynthetic parameters remained unaltered (**Table 1**). Light response curves of qN and qL did not reveal any major differences to the wild type (**Figure 7A**). Likely, due to limitation of PSII biogenesis by another so far unknown factor, the additionally synthesized D2 and CP43 are condemned to degradation.

These data also imply that no positive feed-forward regulation of PSII biogenesis by the CES process exists in tobacco. In *C. reinhardtii*, D1 translation is activated in the presence of the D2-cyt b_559_ subcomplex, and the availability of D1 (or more likely the formation of the RC-like complex), in turn, stimulates translation of CP47 by allowing its assembly into the nascent PSII complex (Minai et al., 2006). In tobacco, despite the increased transcript abundance and translation of *psbD, psbC*, and *psbZ*, the transcript abundance and translation of both *psbA* and *psbB* remained unaltered. Also, for the other plastome-encoded subunits of PSII, no obvious changes in transcript abundance or translation were observed in response to the increased transcription and translation of *psbD* and *psbC*, except for a slightly reduced translation of *psbK* (see below). Therefore, increased synthesis of D2 does not activate the translation of the downstream subunits by sequestering free, unbound proteins.

Another important conclusion from the analysis of the *s-aadA-psbD* mutant is that, in tobacco, translation of *psbD* and *psbC* is either independent of mRNA-specific translation initiation factors, or these are present in large excess. If any such factor were limiting, overexpression of the transcript would have resulted in lower translation efficiency. Chloroplast mRNAs with an SD in their 5’-UTR may be less dependent on mRNA-specific translation factors than those that do not contain an SD (Scharff et al., 2017; Zoschke and Bock, 2018). Alternatively, because translation of the small PsbK subunit was slightly decreased in the s-*aadA-psbD* and the s-GTG-*psbD* mutant (**Figure 4**), a specific translation factor might be shared between *psbD* (and/or *psbC*) and *psbK*, possibly explaining its decreased translation.

### Translation initiation mutants

In addition to increasing and decreasing the mRNA abundance of the operon genes, we altered *psbD* translation initiation efficiency via the introduction of point mutations into the initiation codon. This approach had been previously applied to reduce accumulation of chloroplast ATP synthase by the introduction of point mutations into the start codon of the catalytic AtpB subunit (Rott et al., 2011), and to repress the essential plastid-encoded ClpP subunit of the stromal Clp protease (Moreno et al., 2017). While in the case of the chloroplast ATP synthase, the *GTG-atpB* mutation had a more severe effect on ATP synthase accumulation than the *TTG-atpB* mutation (Rott et al., 2011), in the case of ClpP, the *TTG-clpP* mutation resulted in a more severe defect (Moreno et al., 2017). Here, ribosome footprint analysis (**Figure 4**) revealed that the GUG initiation codon was almost as efficient as the AUG start codon for *psbD* (**Supplemental Figure 1**). Translation efficiency of *psbD* in the s-TTG-*psbD* mutant was strongly reduced, showing that the UUG initiation codon is less efficient than either AUG or GUG in the sequence context of the *psbD* 5’-UTR (**Supplemental Figure 1**). Accordingly, different from all other s-mutants, the s-TTG-*psbD* mutant suffered from a mild reduction of PSII content (**Table 1**). Furthermore, the as-GTG-*psbD* mutant was viable under autotrophic conditions, while the as-TTG-*psbD* mutant was not, again suggesting that translation initiation from the TTG codon was less efficient. Finally, despite the strongly diminished translation efficiency of *psbD* in the s-TTG-*psbD* mutant, translation of *psbC* remained unaffected, arguing against a strong coupling of *psbC* translation initiation and termination of *psbD* protein synthesis, which had been previously suggested based on *in vitro* studies (Adachi et al., 2012).

We also generated mutants with an altered SD, because *psbD* translation was severely affected in transplastomic tobacco plants harboring a mutation in the anti-SD (Scharff et al., 2017). Overall, the behavior of the SD mutants was very similar to the start codon mutants. The s-SD-*psbD* mutants were similar to the s-GTG-*psbD* mutant, in that no differences to the wild type were observable (**Table 1, Figures 5-7**).The as-GGAGCA-*psbD* mutant was most similar to as-GTG-*psbD*, while the other two as-SD-*psbD* mutants did not survive under autotrophic conditions, similar to the as-TTG-*psbD* mutant.

### D2 and the PSII repair process

All three as-mutants, which were viable under autotrophic conditions, were light-sensitive and had to be grown under a reduced light intensity of 100 μE m^-2^ s^-1^. Despite this reduced actinic light intensity, their growth was strongly retarded (**Figure 5**), and they suffered from severe photosynthetic defects (**Table 2, Figure 7B,D**). The reduced maximum quantum efficiency of PSII in the dark-adapted state is at least partly attributable to the presence of uncoupled LHCII in the as-mutants (**Figure 7D**). Our data suggest that the diminished synthesis of D2 renders the mutants more sensitive to photoinhibition. Normally, the D1 protein is most prone to photodamage. Damaged D1 is replaced by the PSII repair cycle, which depends on PSII core protein phosphorylation, dissociation of the PSII core from its LHCII, and its diffusion from the grana into unstacked thylakoids. There, the damaged D1 is co-translationally replaced by a newly synthesized D1 protein (Herbstová et al. 2012, Yamamoto et al., 2014; Järvi et al. 2015). Using ribosome profiling, Chotewutmontri and Barkan (2018) and Schuster et al. (2020) showed that, while translation initiation of *psbA* is activated in response to light stress, translation of other plastid-encoded PSII subunits remains unaltered.

Our observation that the low amount of PSII accumulating in the *as-aadA-psbD* mutant is prone to photoinhibition is difficult to explain with a scenario, where exclusively D1 is damaged. The lowered total PSII accumulation in these mutants should reduce photoinhibition, as less electrons are fed into the electron transport chain, thus likely reducing the rate of reactive oxygen species formation. However, an increased turnover of both D2 and PsbH, relative to other subunits of PSII, has been reported (Rokka et al., 2005). Recently, higher protein turnover rates of D2 and CP43 than of most other PSII subunits have been determined; however, D1 still had a far higher turnover rate than D2 (Li et al., 2017). Our mutants will provide useful tools for future studies of the repair of PSII accumulating damaged D2 protein due to reduced synthesis levels.

### What limits PSII biogenesis in vascular plants?

Because our data show that neither D2 nor CP43 limits PSII accumulation in tobacco, the question remains, what ultimately restricts PSII accumulation. A limiting function of one of the other plastome-encoded subunits of PSII appears unlikely, given that a rate-limiting step should occur early in the pathway. The next subunits to assemble into the D2-cyt b_559_ subcomplex are D1 and the small PsbI subunit. PsbI is not essential for PSII accumulation (Schwenkert et al., 2006). D1 is an unlikely candidate to control PSII biogenesis, because its rapid turn-over would generate the problem to separately regulate its the synthesis for the PSII repair cycle and for *de novo* assembly of PSII. All other subunits are incorporated at a later point into nascent PSII. If they would limit PSII biogenesis, this would not only be wasteful, but also bear the risk of increased production of reactive oxygen species, because the RC-like complex is already capable of charge separation at P_680_. PSII assembly intermediates, that are already capable of charge separation, but do not yet contain fully functional donor and acceptor sides, are more prone to photooxidative damage than mature PSII (Mamedov and Styring, 2003; Shevela et al., 2019). Therefore, the highly regulated rate-limiting step of PSII biogenesis should occur prior to the formation of these dangerous reactive assembly intermediates, to avoid unnecessary photodamage.

While a limiting function of auxiliary proteins may seem to be a rather indirect regulatory mechanism, the plastid-encoded assembly chaperone Ycf3 can restrict PSI biogenesis (Petersen et al., 2010; Schöttler et al., 2011). The ultimate proof that a subunit or an auxiliary protein indeed limits PSII biogenesis can only be provided by establishing a strict correlation between the its content and PSII accumulation over a wide concentration range, including the demonstration that overexpression of the factor leads to overaccumulation of functional PSII, as reported for *C. reinhardtii* (Klinkert et al., 2006). Also, such a correlation should be validated under different growth conditions. For example, overexpression of the *petA* mRNA increased cyt b_6_f accumulation only in low-light conditions. In high light or under adverse growth conditions such as heat or chilling stress, no difference in cyt b_6_f content between the wild type and the *petA* mRNA overexpressor was discernable anymore (Schöttler et al., 2017).

The identification of the bottleneck in PSII biogenesis may also pave the way to targeted manipulations of PSII content, without interfering with PSII subunit composition and function. The latter has been observed in many mutants with decreased PSII contents due to the repression of individual subunits. Most of these mutants contain less PSII, but the residual PSII is functionally impaired (reviewed by Shi et al., 2012; Plöchinger et al., 2016). By specifically manipulating the rate-limiting step of the assembly process, only fully functional PSII of wild-type structure should accumulate. Such mutants would be excellent tools for basic research on photosynthesis, but also would open up new opportunities in biotechnology and plant breeding.

## Material and methods

### Plant material and growth conditions

Tobacco plants (*Nicotiana tabacum* cv Petit Havana) were grown under aseptic conditions on agar-solidified Murashige and Skoog medium containing 30 g/L sucrose (Murashige and Skoog, 1962). Transplastomic lines were rooted and propagated on the same medium. For seed production, transplastomic plants were grown in soil under standard greenhouse conditions. Inheritance and seedling phenotypes were analyzed by germination of surface-sterilized seeds on Murashige and Skoog medium containing 500 mg/L spectinomycin. For photosynthesis measurements and molecular analyses, seeds of tobacco wild-type and transplastomic plants were germinated under long-day conditions (16 h light) at a light intensity of 100 μE m^-2^ s^-1^. The day temperature was 22°C, the relative humidity 75%. During the night, temperature and relative humidity were decreased to 18°C and 70%, respectively. Four weeks after germination or later (in the case of mutants severely retarded in growth, see results), plants were transferred to a Conviron chamber with 350 μE m^-2^ s^-1^ light intensity. All other environmental parameters were kept constant. After a minimum time of 14 days to allow full acclimation to the new environment, all measurements were performed on the youngest fully expanded leaves at the onset of flowering.

### Vector construction for plastid transformation

The region of the tobacco plastid genome containing a part of the *psbD* gene was isolated as a 1.5-kbp KpnI/SacI fragment (**Figure 1A**) with primers PsbD_F and PsbD_R (**Table S1**), and then cloned into the pBC SK+ plasmid vector (Stratagene). The resulting construct was used to produce the translation initiation codon mutations by PCR. The region surrounding the SD and the ATG start codon was cleaved with BsaHI and EcoO109I (**Figure 1A**) and replaced by PCR products carrying the point mutations. The mutations of the SD or the start codon were introduced with one of the 5 primers with mutated sites (PsbD_mut1_F to PsbD_mut5_F) and primer PsbD_mut_R as the reverse primer for all the PCR amplifications (**Table S1**). A chimeric *aadA* gene fused to chloroplast-specific expression signals and conferring resistance to the aminoglycoside antibiotics spectinomycin and streptomycin (Svab and Maliga, 1993) was cloned into a unique NdeI site within the 5’-UTR of the *psbD* gene (**Figure 1A**) to enable selection of transplastomic lines. Digestion of the final transformation vectors with restriction enzyme KpnI was used to identify the orientations of *aadA* insertion. Both the sense and the antisense orientation of the *aadA* relative to *psbD* were used for plastid transformation.

### Generation of transplastomic plants

Young sterile leaves of tobacco cultivar Petit Havana were bombarded with the respective transformation vectors bound to gold particles (0.6-μm diameter) using a helium-driven biolistic gun (PDS-1000 He; Bio-Rad). Primary transformants were selected from 5 × 5-mm leaf pieces exposed to plant regeneration medium containing 500 mg/L spectinomycin. Several independent transplastomic lines were then subjected to a maximum of three additional rounds of regeneration on spectinomycin-containing medium to enrich the transplastome and select against residual copies of the wild-type plastome. Spontaneous spectinomycin-resistant plants were eliminated by a double resistance test on medium supplemented with 500 mg/L spectinomycin and 500 mg/L streptomycin (Svab and Maliga, 1993; Bock, 2001; Bock 2015). Homoplasmic transplastomic lines were transferred to the greenhouse for seed production. The homoplasmic state of the progeny was confirmed by inheritance test and RFLP analysis (**Figure 1B,C)**. The presence of the mutations was confirmed by DNA sequencing. It should be noted that the as-*aadA-psbD* mutant had been previously used as a control by Krech et al. (2013).

### Isolation of nucleic acids and hybridization procedures

Total plant DNA was isolated from fresh leaf material by a rapid cetyltrimethylammoniumbromide-based miniprep procedure (Doyle and Doyle, 1990). DNA samples digested with restriction enzymes were separated in 1% agarose gels and blotted onto Hybond-XL nylon membranes (GE Healthcare) according to the manufacturer’s instructions. RNA was extracted using the peqGold TriFast reagent (Peqlab). RNA samples were separated in 1% formaldehyde-containing agarose gels and blotted onto Hybond-XL nylon membranes. Hybridization probes were generated by PCR amplification using specific oligonucleotides (**Table S1**). Prior to labeling, DNA fragments were purified by agarose gel electrophoresis followed by extraction from an excised gel slice using a NucleoSpin Extract II kit (Macherey-Nagel). Hybridization probes were labeled with α[^32^P]dCTP by random priming (Multiprime DNA labeling kit; GE Healthcare). Hybridizations were performed at 65°C in Rapid-Hyb buffer (GE Healthcare) according to the manufacturer’s instructions. Hybridization signals were quantified using a Typhoon Trio+ variable mode image (Amersham Biosciences) and Image Quant 5.2 software.

### Polysome loading assays

Isolation of polysomes and RNA extraction from sucrose gradient fractions were performed as described previously (Barkan, 1998; Rott et al., 2011). Equal aliquots of extracted RNAs from each fraction were separated by denaturing agarose gel electrophoresis as described above. For the puromycin control, polysome samples with 0.5 mg/mL puromycin were incubated at 37°C for 10 min prior to ultracentrifugation.

### Ribosome profiling

Ribosome footprints and total RNA were isolated as described in Zoschke et al. (2013), except that after micrococcal nuclease treatment to degrade nuclease-accessible mRNA and cleave polysomes into monosomes, 4 ml lysate was layered onto a 1 ml sucrose cushion (30% (w/v) sucrose, 0.1 M KCl, 40 mM Tris-acetate, pH 8.0, 15 mM MgCl_2_, 5 mM 2-mercaptoethanol, 100 μg/ml chloramphenicol, and 100 μg/ml cycloheximide) and centrifuged for 1.5 h at 50,000 rpm at 4 °C in a SW55 Ti rotor (Beckman). RNA labeling was performed according to Zoschke et al. (2013) with the following minor modification: 4 μg purified footprints and 3.5 μg fragmented total RNA derived from mutant and wild-type plants were differentially labeled with Cy3 and Cy5 (ULS Small RNA Labeling Kit, Kreatech Diagnostics), respectively, following the manufacturer’s instructions.

Ribosome and transcriptome profiling data were analyzed as described in Trösch et al. (2018). Briefly, all local background-subtracted single-channel signals (F635-B635 and F532-B532, respectively) were normalized to the average signal of the datasets of the three mutants and their corresponding wild type including all replicates of ribosome footprints and total mRNA to remove biases introduced by technical variation. The average of the probe signals for each reading frame (RF). Then, the average values were log_2_-transformed. The relative abundance of ribosome footprints and total mRNA were calculated for each RF by normalization of the average of each RF to the average signal of all RFs. By doing so we obtained for each RF relative expression levels (RNA or ribosome footprint), which show expression of the specific RF in relation to the average of all chloroplast reading frames. Translation efficiencies were calculated for each RF by subtracting the summarized log_2_-transformed signals of the total mRNA from the summarized log_2_-transformed signals of ribosome footprints. The average and standard deviation of relative abundances of ribosome footprints, total mRNA, and translation efficiency were calculated for each RF from three biological replicates.

### Chlorophyll-a fluorescence measurements and leaf absorptance

A F-6500 fluorometer (Jasco Inc., Groß-Umstadt, Germany) was used to measure 77 K chlorophyll-a fluorescence emission spectra on freshly isolated thylakoid membranes equivalent to 10 μg chlorophyll mL^-1^. The sample was excited at 430-nm wavelength with a bandwidth of 10 nm, and the emission spectrum was recorded between 655- and 800-nm in 0.5-nm intervals with a bandwidth of 1 nm. The spectra were normalized to the PSI emission maximum at 734 nm, or to the maximum emission of PSI-LHCI in the as-mutants with shifted emission maximum.

*In vivo* measurements of chlorophyll-a fluorescence parameters at 22°C were performed using the modular transmittance version of the Dual-PAM (Heinz Walz GmbH). Light-response curves of linear electron flux, non-photochemical quenching (qN; Krause and Weis, 1991), and the redox state of the PSII acceptor side (qL; Kramer et al., 2004) were measured after 30 min of dark adaptation. The light intensity was increased stepwise from 0 to 2500 μE m^-2^ s^-1^, with a measuring time of 150 s for each light intensity under light-limited conditions and of 60 s under light-saturated conditions. Linear electron transport was corrected for leaf absorptance, which was calculated from leaf transmittance and reflectance spectra as 100% minus transmittance (%) minus reflectance (%). Spectra were measured between 400- and 700-nm wavelengths using an integrating sphere attached to a photometer (V-550, Jasco Inc.). The spectral bandwidth was set to 1 nm, and the scanning speed was 200 nm min^-1^.

### Thylakoid membrane isolation and photosynthetic complex quantification

Thylakoid membranes were isolated as described previously (Schöttler et al., 2004). The chlorophyll content and a/b ratio were determined in 80% (v/v) acetone according to Porra et al. (1989). The contents of PSII and the cyt b_6_f were determined from difference absorbance signals of cyt b_559_ (PSII) and the cytochromes b6 and f in destacked thylakoids equivalent to 50 μg chlorophyll mL^-1^ (Kirchhoff et al., 2002). All cytochromes were fully oxidized by the addition of 1 mM potassium hexacyanoferrate (III). Then, 10 mM sodium ascorbate was added to reduce the high-potential form of cyt b_559_ and cytochrome f. Finally, the addition of 10 mM sodium dithionite reduced the low potential form of cyt b_559_ and the two b-type hemes of cytochrome b6. Using a V-550 spectrophotometer equipped with a head-on photomultiplier (Jasco GmbH) at each of the three redox potentials, absorbance spectra were measured between 575- and 540-nm wavelengths. The spectral bandwidth was 1 nm and the scanning speed 100 nm min^-1^. Ten spectra were averaged per redox condition. Difference spectra were calculated by subtracting the spectrum measured in the presence of hexacyanoferrate from the ascorbate spectrum, and by subtracting the ascorbate spectrum from the spectrum measured in the presence of dithionite, respectively. Finally, a baseline calculated at wavelengths between 540 and 575 nm was subtracted from the signals. Then, the difference spectra were deconvoluted using reference spectra as previously described (Kirchhoff et al., 2002; Lamkemeyer et al., 2006). PSI was quantified from light-induced difference absorbance changes of P700. Thylakoids equivalent to 50 μg chlorophyll mL^-1^ were solubilized in the presence of 0.2% (w/v) n-dodecyl-β-D-maltoside (DDM). After the addition of 10 mM sodium ascorbate as the electron donor and 100 μM methylviologen as the electron acceptor, P700 photo-oxidation was achieved by applying a light pulse of 250 ms (2000 μmol photons m^-2^ s^-1^). Measurements were performed with the Pc-P_700_ version of the Dual-PAM instrument (Heinz Walz GmbH). Pc contents, relative to PSI, were determined in intact leaves and then recalculated based on the absolute PSI quantification performed in isolated thylakoids (Schöttler et al., 2007).

### Protein gel electrophoresis and immunoblotting

Thylakoid proteins were separated by SDS-PAGE and then transferred to a polyvinylidene membrane (Hybond P, GE Healthcare) using a tank blot system (Perfect Blue Web M, VWR International GmbH, Darmstadt, Germany). Immunochemical detection was performed using an enhanced chemiluminescence detection reagent (ECL Prime, GE Healthcare) according to the manufacturer’s instructions. Chemiluminescence was detected on X-ray film. Antibodies against the photosynthetic proteins were purchased from Agrisera AB (Vännäs, Sveden).

## Acknowledgements

We are grateful to Stefanie Seeger and Claudia Hasse (MPI-MP) for help with chloroplast transformation. We thank the MPI-MP Green Team for plant care and cultivation.

## Supplemental Data

**Supplemental Table S1.** Summary of oligonucleotide sequences.

**Supplemental Figure S1. Ratios of relative average translation output and transcript accumulation levels.**

**Supplemental Figure S2. Reproducibility of transcript abundance and ribosome footprint data between biological replicates.**

